# The evolutionary advantage of cultural memory on heterogeneous contact networks

**DOI:** 10.1101/466524

**Authors:** Oana Carja, Nicole Creanza

## Abstract

Cultural processes, as well as the selection pressures experienced by individuals in a population over time and space, are fundamentally stochastic. Phenotypic variability, together with imperfect phenotypic transmission between parents and offspring, has been previously shown to play an important role in evolutionary rescue and (epi)genetic adaptation of populations to fluctuating temporal environmental pressures. This type of evolutionary bet-hedging does not confer a direct benefit to a single individual, but instead increases the adaptability of the whole lineage.

Here we develop a population-genetic model to explore cultural response strategies to temporally changing selection, as well as the role of local population structure, as exemplified by heterogeneity in the contact network between individuals, in shaping evolutionary dynamics. We use this model to study the evolutionary advantage of cultural bet-hedging, modeling the evolution of a variable cultural trait starting from one copy in a population of individuals with a fixed cultural strategy. We find that the probability of fixation of a cultural bet-hedger is a non-monotonic function of the probability of cultural memory between generations. Moreover, this probability increases for networks of higher mean degree but decreases with increasing heterogeneity of the contact network, tilting the balance of forces towards drift and against selection.

These results shed light on the interplay of temporal and spatial stochasticity in shaping cultural evolutionary dynamics and suggest that partly-heritable cultural phenotypic variability may constitute an important evolutionary bet-hedging strategy in response to changing selection pressures.

## Introduction

Many foundational models of social learning and cultural evolution are constructed within the framework of theoretical population genetics (Cavalli-Sforza and Feldman, 1981, 1973; Feldman and Cavalli-Sforza, 1976; Cavalli-Sforza et al., 1982; Feldman and Cavalli-Sforza, 1984). With genetic evolution as a starting point, models of cultural evolution emphasize that cultural traits-learned behaviors such as beliefs, practices, and tools—can be transmitted between individuals and are subject to evolutionary forces such as selection and drift (Cavalli-Sforza and Feldman, 1973; Creanza et al., 2012, 2017a). In addition, these population-genetic modeling frameworks facilitate the joint consideration of genetic and cultural traits, allowing researchers to track allele frequencies and cultural phenotypes within the same population and assess their evolutionary effects on one another (Feldman and Zhivotovsky, 1992; Laland et al., 1995, 2000; Odling-Smee et al., 2003).

These models of cultural evolution have generally assumed a well-mixed population, meaning that any individual is equally likely to interact with and learn from any other individual in the population. In this scenario, researchers can use the frequency of cultural traits in the population to approximate the probabilities that individuals with those traits will interact with one another. However, humans often bias their interactions based on perceived cultural similarities to other individuals, showing preferences for learning from other individuals that are more similar to them in certain ways. This phenomenon, known as homophily or assortment, implies that rare traits in the population might be transmitted more often than predicted by their low frequency if the people with those traits preferentially interact with one another (Creanza and Feldman, 2014; Centola, 2010, 2011). In this sense, homophily acts to increase the perceived frequency of a trait in a subset of the population, even if its frequency is low in the population as a whole. In addition, homophily might bias the connections in a network, if links are formed between individuals who share certain existing beliefs and behaviors (Centola, 2011; McPherson et al., 2001).

Even in the absence of homophily, human interactions are unlikely to be ideally represented by a well-mixed population. Humans have complex contact networks, where interactions between some individuals are common and interactions between other individuals are rare or absent (Ohtsuki et al., 2006; Christakis and Fowler, 2008, 2013). Some of these differences in interaction might be due to the spatial distribution of individuals in a population; individuals located geographically more close to one another are more likely to interact. Other differences might be driven by social structure, with interactions on a social network more likely to occur between genetically related individuals and between individuals sharing social contexts (Henrich and Broesch, 2011; Apicella et al., 2012).

Taken together, these studies hint at the potentially important role of local population structure in human cultural transmission and the necessity of theoretical treatments of cultural evolution that account for complex contact networks. However, few models explore how spatial or network structure can affect the spread of a cultural trait compared to well-mixed populations. Cavalli-Sforza and Feldman (1981) used reaction-diffusion dynamics to characterize the spread of information in a population, making the analogy between the spatial interactions of individuals and those of molecules in a chemical reaction. Derex and Boyd (2016) used theoretical and empirical research to demonstrate that certain types of network structure might facilitate human innovations: when people could observe all other members of a population attempting to solve a problem, the population often became “stuck” on a good solution to a problem but did not find the best possible solution. On the contrary, if subsets of the population worked separately and then compared their solutions, this partially connected network produced a diversity of perspectives that often produced better combined results than completely connected populations (Derex and Boyd, 2016). Creanza et al. (2017b) analyzed spatially structured populations from another perspective, modeling a set of separate populations with their own cultural repertoires and allowing these populations to interact through migration. These migration events between populations produced bursts of innovations that increased the cultural repertoires of populations that received migrants. Separately, it has been shown that network structure can act both as an amplifier and a suppressor of selection (Lieberman et al., 2005; Hindersin and Traulsen, 2015), underscoring the potential relevance of contact structure in shaping the evolution of cultural traits.

In addition to homophily and network structure, previous research also suggests that fluctuating environments can shape cultural evolutionary dynamics: some cultural traits—for example, behavioral adaptations to cold weather such as fur clothing—might be more useful in some environments than in others (McCartney, 1975; Gilligan, 2010). One sense in which cultural evolution is fundamentally different from genetic evolution is that environmental pressures can drive the innovation of new cultural traits: humans exposed to cold weather can invent warmer clothing in response, whereas a bacterium exposed to antibiotics cannot invent a resistance mutant in direct response. Thus, cultural evolutionary patterns could differ in harsh environments compared to stable environments, particularly if the harsh environments provide challenges that can be ‘solved’ with new behaviors or technologies (Rendell et al., 2010; Smaldino et al., 2013; Fogarty et al., 2015; Fogarty and Creanza, 2017; Fogarty, 2018).

Patterns of environmental change can also influence learning strategies (Borenstein et al., 2008; Aoki and Feldman, 2014). For example, if the environment fluctuates quickly, individuals are more likely to experience a different environment from their parents, which might decrease the fitness value of high-fidelity vertical (parent-to-offspring) transmission. Cultural evolution models by Fogarty (2018) have also demonstrated that the rate of cultural innovation in a population decreases with environmental stability and increases in unstable, periodically changing environments. These results suggest that, similar to (epi)genetic systems (Carja and Feldman, 2011; Carja et al., 2014a, b), partly heritable cultural trait variation can act as a phenotypic bet-hedging strategy, using dynamic regulation of cultural phenotypic variability to facilitate adaptation to changing environmental pressures. While cultural plasticity can cause phenotypes to differ widely within a cultural lineage, and fixed phenotypes only allow offspring with identical traits to the parents, the type of partly heritable cultural memory we explore here can produce cultural heterogeneity with familial correlations intermediate to these two extremes. In this context, cultural memory represents a different phenomenon than two related parameters that are often discussed in the cultural evolution literature: the cultural transmission probability, the likelihood that another individual adopts one’s traits, and the transmission fidelity, the likelihood that the other individual accurately reproduces the cultural trait that it was attempting to copy. Here, cultural memory applies to general traits that can take one of multiple specific forms: a child of an agriculturalist might be exceedingly likely to also become an agriculturalist, but the level of cultural memory would influence whether the child chooses to farm the specific crop of his parent versus choosing a different crop used by a neighbor, particularly when the neighbor appears to have an excess of food. With higher levels of cultural memory, the child is more likely to adopt the parent’s specific form of the trait.

In sum, understanding the spatial and environmental dynamics of human cultural interactions could shed light on some of the fundamental aspects of human behavior, such as social learning, cooperation, and cumulative culture. Our paper studies cultural strategies in temporally varying environments by analyzing the evolutionary advantage of partly heritable cultural memory, and how this evolutionary advantage is shaped both by fluctuating environments and local population structure. It starts by specifying a Moran death-birth model of cultural transmission for spatially structured populations, subject to periodic environmental change. Here, the spatial component is represented by a heterogeneous contact network between the individuals in the population: each node represents an individual, and the interactions between individuals are represented by connections between the nodes. The *degree* of a node is the number of its connections to other nodes (Wasserman and Faust, 1994); it follows that the mean degree of a network is the average number of connections across all nodes. When the degree distribution of the nodes of a network has a small standard deviation, the nodes of the network are generally similar in degree; however, when the standard deviation in degree is large, nodes with few or many connections are more common (**Figure 1**).

**Figure 1:**
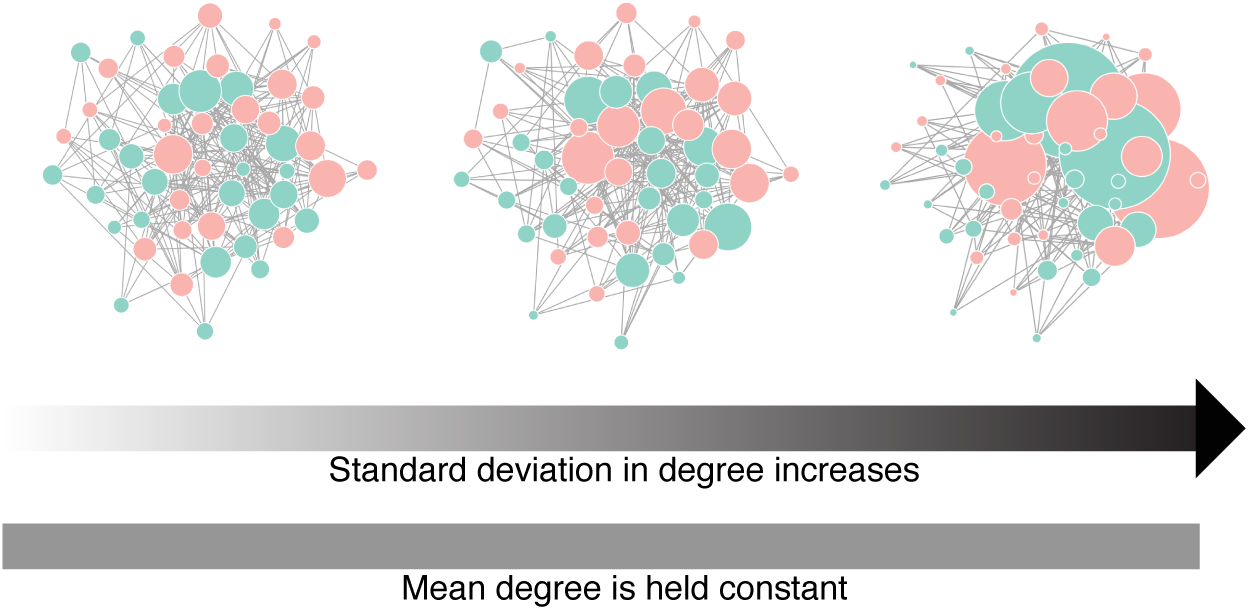
Networks that differ by standard deviation in degree. The three networks have the same mean degree; in other words, the nodes in each network have the same mean number of connections. However, they differ in their standard deviation in degree: networks with high variance in degree have many small-degree nodes and some large-degree nodes, also referred to as hubs. Here, the size of a node corresponds to its degree and node color is arbitrary.

We pose our research question in terms of analyzing the probability of fixation of a mutant cultural trait that permits a range of expressible, partly heritable cultural phenotypes. This model builds on previous cultural evolutionary theory by incorporating essential, often neglected aspects of cultural transmission: spatial structure of transmission networks and partially heritable phenotypic variability (building on Carja and Plotkin (2017)). Similar to Carja and Plotkin (2017), we find that the evolutionary advantage of a phenotypically variable cultural trait critically depends on the cultural memory of individuals expressing the variable trait. When the population experiences multiple environmental changes, the probability of fixation of a phenotypically variable trait depends non-monotonically on the probability of cultural memory. We also show that the population structure of interactions on a social network suppresses the probability of fixation of such a trait, compared to a well-mixed population. Further, this suppression is stronger for networks of increased standard deviation in degree. We provide intuition for the complex dependence of this evolutionary advantage on the degree of phenotypic memory and the heterogeneity of the contact network, and we discuss the interacting roles of population structure, environmental change, and phenotypic memory on the evolution of cultural variability.

## Model

We use a death-birth Moran-type model to describe changes in the frequency of a cultural ‘allele’ in a finite population of fixed size *N*. Each individual is defined by a single biallelic cultural locus *A/a*, which controls its phenotypic range. Each individual has one allele, *A* or *a*, and there is uniparental inheritance. Initially, we assume that all individuals have the non-variable *A* allele, with one individual with the phenotypically variable *a* allele introduced at time *t* = 0. The *A* allele encodes a fixed phenotypic value, whereas individuals with the *a* allele may express a wider range of phenotypes. Here, the phenotypic distribution of the *a* allele is chosen from a discrete uniform distribution with probability mass on two points. In other words, the phenotype of individuals with the *a* allele can take one of two forms (**Figure 2**). In previous work, it was shown that this representation did not qualitatively differ from a scenario in which the variable phenotype was drawn from a continuous uniform distribution (Carja and Plotkin, 2016).

**Figure 2:**
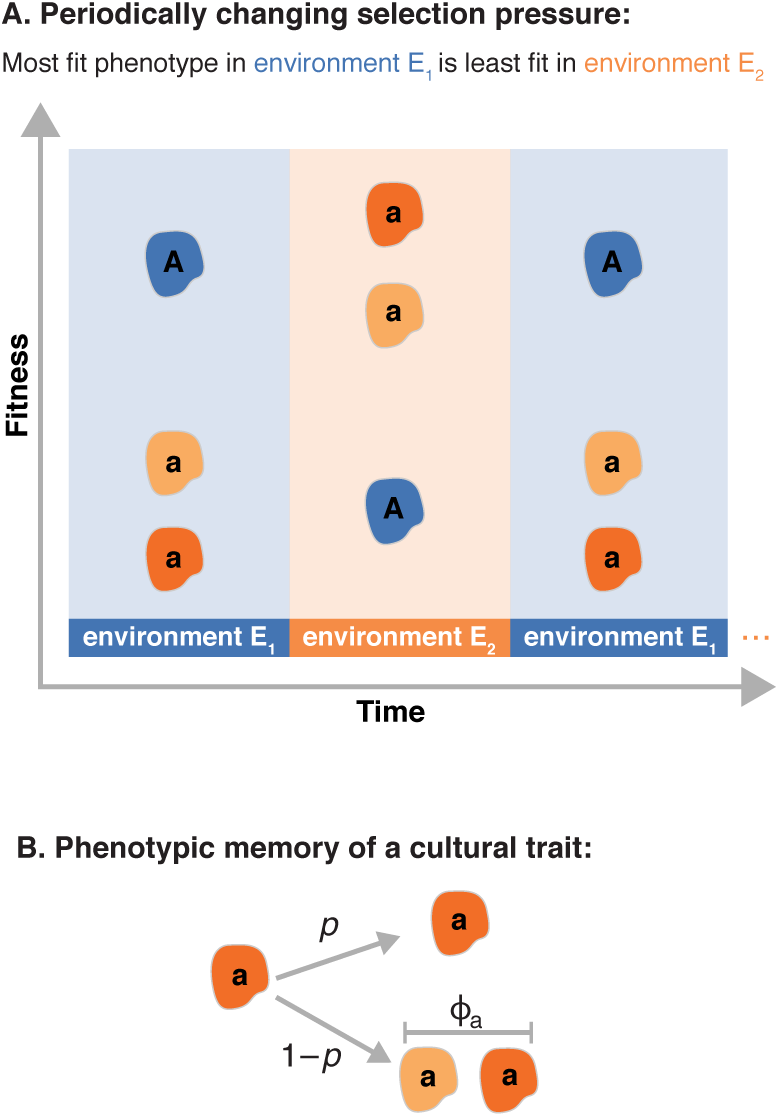
Illustration of the model. **Panel A:** Periodically changing selection pressures. Time is shown on the *x*-axis and fitness on the *y*-axis. Every *n* time steps, the environment changes, alternating between *E*_1_ and *E*_2_. The phenotype with the highest fitness in *E*_1_ has the lowest fitness in *E*_2_ and vice versa. The mean fitness of the *a* phenotypes in one environment equals the fitness of the *A* phenotype in the other environment. **Panel B:** Phenotypic memory of a cultural trait. When an individual *a* gives birth, with probability *p* (the probability of phenotypic memory), its offspring inherits the cultural trait of its parent, and with probability 1 – *p*, the offspring’s phenotype is resampled from the phenotypic distribution. Adapted from Carja and Plotkin (2017).

Since the phenotype of an individual translates to its fitness, this system could usefully be applied to many types of cultural traits. Here, we use a hunting and gathering example to illustrate a cultural system that is influenced by environmental change, social network connections, and phenotypic cultural memory. Say that the *A* allele represents a generalist cultural phenotype of gathering food from the environment. The *a* allele could then be a cultural phenotype of hunting animals for food, with the added difference that there are multiple forms of the *a* allele, each representing different hunting specializations, including tools or techniques that are culturally transmitted. For example, individuals with the *A* allele have the cultural phenotype of gathering food, and individuals with the *a* allele have one of multiple cultural phenotypes with different fitness values, say, hunting either small game with traps or hunting large game with spears. Individuals with the *a* allele specialize in one type of hunting and the tools associated with it. For this modeling application, we make the simplifying assumption that fitness is closely linked to an individual’s food sources, which might not generalize to populations with the cultural practice of sharing all of their food with the group.

We introduce two environments, *E*_1_ and *E*_2_. In these environments, individuals of the wild type *A* and mutant type *a* each give birth and die according to the following per-capita rates:

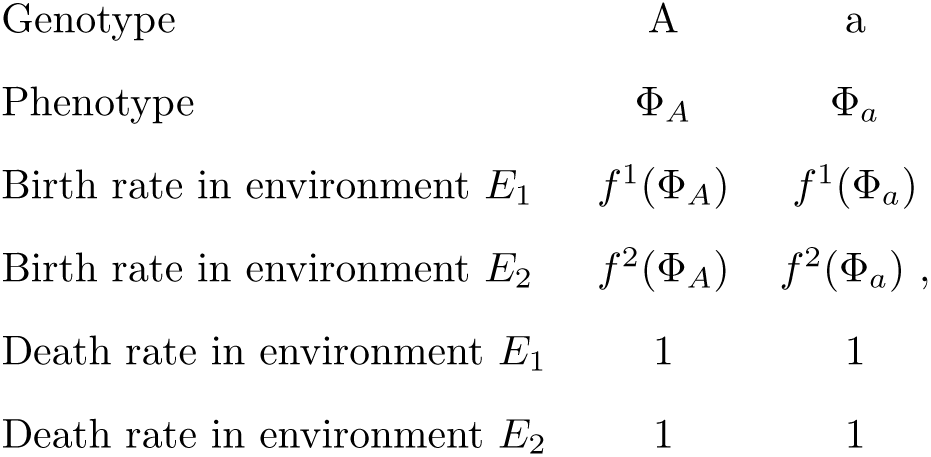

where Φ_*a*_ denotes a random variable, and Φ_*A*_ is a fixed value. The functions f^*i*^ : ℝ → ℝ (*i* ∈ {1, 2}) map phenotype to birth rate in each of the two environments, and *f*^1^ is the identity function. We assume that both alleles have the same expected mean fitness in their optimal environment, and the same expected mean fitness in their unfavorable environment: 𝔼(*f*^1^(Φ_*A*_)) = 𝔼(*f*^2^(Φ_*A*_)) and 𝔼(*f*^2^(Φ_*A*_)) = 𝔼(*f*^1^(Φ_*a*_)). This condition also ensures that the average of two alleles’ mean fitnesses, which we denote 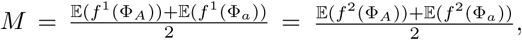 is the same in both environments. The function *f*^2^ is defined as a reflection of *f*^1^ around *M*: *f*^2^(*x*) = 2*M* – *f*^1^(*x*). As a result, the variance in fitness of allele *a* with randomly drawn phenotype is the same in both environments: Var(*f*^1^(Φ_*a*_))=Var(*f*^2^(Φ_*a*_)) =Var(Φ_*a*_) (as in Carja and Plotkin (2017)).

In this model, there are also multiple epochs of environmental changes, occurring periodically. The mapping from phenotype to fitness depends on the environmental regime, and it is chosen so that both alleles have the same expected fitness across environments, so that the only difference between them is the possibility of (partly heritable) phenotypic variability. We choose phenotypic ranges and fitness functions so that the mean fitness expressed by each of the two genotypes are equal when averaged over the two environmental regimes. This setup allows us to focus on the evolutionary advantage of the phenotypic variance of *a*, Var(Φ_*a*_), and to study population persistence without conflating this effect with any mean-fitness advantage. In our analysis of periodic environmental changes, we assume that the population experiences two different types of environments, *E*_1_ and *E*_2_, which alternate deterministically every *n* time steps, so that both environments are experienced every 2*n* time steps. We assume that one environment is more favorable to one allele, and the other environment to the other allele – that is, we study a model where the phenotypically variable allele *a* has lower expected fitness than the wild-type allele in one of the environmental regimes, and it has higher expected fitness than the wild-type in the other regime (as illustrated in **Figure 2**). In other words, the cultural trait with highest fitness in the first environment has the lowest fitness in the second environment, and the trait with the lowest fitness in the first environment will have the highest fitness in the second environment.

In the context of our hunting and gathering example, this scenario represents the notion that the reproductive fitness of individuals who gather different food sources can differ based on aspects of the environment, including differences in environment due to the population migrating to a new area. For example, during a high-rainfall environmental epoch, certain plants might grow well and be rich food sources for foragers, whereas during a low-rainfall epoch, the same types of plants have much lower yield. In contrast, perhaps it is easier to find and catch prey when foliage is less dense during a dry epoch, whereas dense foliage increases the difficulty of hunting and lowers yields. Thus, the fitness of individuals with a stable preference for a certain food type would change over time as the environment fluctuates. We can envision a similar fluctuation in hunting and gathering success if the population is nomadic and migrates through wetter and drier environments.

The model essentially differs from Carja and Plotkin (2017) by the introduction of population structure: individuals are nodes of a graph, with links between them representing the contact structure. We generated networks for these simulations using the *igraph* package and the Barabasi-Albert model of preferential attachment. Each network starts with a single vertex and no edges, and nodes are added to reach population size *N*. Each new node is added to the network and connected to other individuals, each with a probability proportional to the individual’s current degree. This family of graphs allows us to easily vary the mean degree or the standard deviation in degree, while keeping the other constant, by varying the power of preferential attachment of the graphs.

Once the network is generated, it remains fixed. We begin at time *t* = 0 with a population in which every node of the network contains individuals fixed for the non-variable *A* allele; on one randomly selected node, we introduce a phenotypically variable *a* allele. At every time step of the Moran model, one individual is chosen to die and a neighbor is chosen, with probability proportional to fitness in the current environment, to replace the empty node with an offspring. The network imposes population structure in the sense that any individual can only pass its *A/a* allele to another node if it is directly connected to that node. However, the connections between nodes do not differ in strength or distance: we consider all connections to be uniform, and an individual’s likelihood of passing on its allele to a connected node is proportional to its fitness in that environment. This is different from a well-mixed population, in which the new birth is chosen proportional to fitness among the entire population.

When a birth occurs, we determine the phenotypic state of the offspring as follows (**Figure 2**). If the individual chosen to reproduce has genotype *A*, then the phenotypic state of the offspring always equals its parent’s (fixed) phenotypic value. For a reproducing individual with the *a* allele, however, there exists a probability of phenotypic cultural memory, denoted by the parameter *p*, between parent and offspring: with probability *p* the offspring retains the phenotypic cultural state of its parent, and with probability 1 – *p* the offspring’s phenotype is redrawn independently from the random variable Φ_*a*_. Thus, individuals of type *a* can express a range of cultural values, and their phenotype is partly heritable between time steps (provided *p* > 0). For our hunting and gathering example, this phenotypic memory represents the likelihood that an *a* offspring will copy the specific hunting tools and techniques of its parent (*p*) rather than sample the distribution of the *a* phenotypes, representing the food sources of the *a* individuals in the population. In the case of periodic environments, we implement environmental changes (and re-calculate event rates) at deterministic times: *n*, 2*n*, 3*n*, etc.

Depending on the timescale considered, this model can apply to multiple forms of cultural evolution. On between-generation timescales, we can consider the death-birth process to represent human reproduction, and the modeled process of cultural learning can represent parent-to-offspring (vertical) transmission of the cultural trait. On within-generation timescales, we can treat the death-birth process to represent a period of cultural sensitivity during which their previous cultural trait can be replaced based on a neighboring individual’s phenotype. This is akin to horizontal cultural transmission, or learning a new cultural trait from one’s peers. Thus, depending on the characteristics of the cultural trait and the relevant timescales represented by each time step, our model can usefully be applied to both vertical and horizontal cultural transmission of a trait on a network. The results of these two conceptions of the model might not be directly comparable with one another since the timescale considered is very different in relation to a human lifespan. In addition to the timescale, the parameters of the model can be tuned to apply to horizontal transmission; for example, the strength of cultural memory might be expected to be lower in horizontal transmission than in vertical transmission, based on empirical studies (Cavalli-Sforza et al., 1982).

With this model, we study the possible long-term advantage of heritable phenotypic variability by analyzing the ability of new phenotypically variable mutations *a* to invade an otherwise non-variable population (*A*) situated on these networks. We define the fixation probability as the proportion of simulations in which the new phenotypically variable allele *a* invades and drives the resident allele to extinction and study how this probability depends on the phenotypic variance, the environmental rate of change, the network structure of the population, and the phenotypic memory associated with the *a* allele. We determine these probabilities of fixation by Monte Carlo simulations, using an ensemble of at least 5,000 replicate populations. Each replicate population undergoes the death-birth process until either the *A* allele or the mutant *a* allele reaches fixation. For all simulations, the population reached fixation in one of the two alleles.

## Results

For a heterogeneous contact network, increasing the number of connections for nodes in the network (in other words, increasing mean node degree) increases the probability of fixation of the *a* allele (**Figure 3**). The heterogeneous contact network acts as a suppressor of selection: the more connected the network, the higher the probabilities of fixation for the *a* allele, with the upper limit being the fixation probability of a well-mixed population.

**Figure 3:**
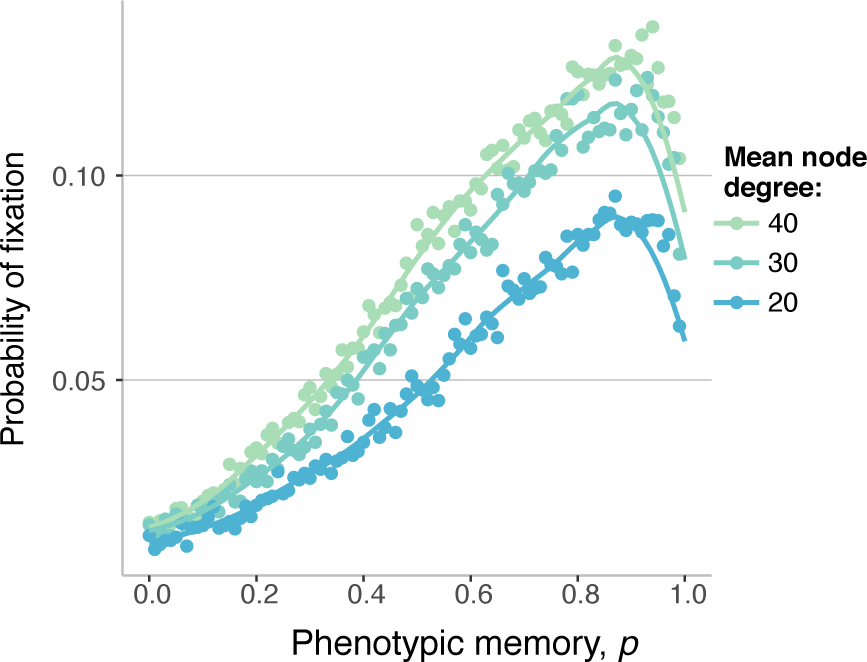
Probability of fixation for different values of the mean degree of the contact network. The lines are splines, while the dots represent the ensemble average across 5000 replicate Monte Carlo simulations. Here *N* = 500, *Φ*_*A*,*E*1_ = 0.8, Φ_*A*,*E*2_ = 0.6, Var(Φ_*a*_)=0.04, the network standard deviation in degree is kept constant at *σ* = 20, and the environment changes every 30 time steps (*n* = 30).

We find that the fixation probability of a phenotypically variable *a* allele is most likely for an intermediate value of the phenotypic memory *p*. We have previously shown that, in well-mixed populations of both fixed and varying size *N*, the probability of fixation of a new plastic allele is a non-monotonic function of the probability of phenotypic memory (Carja and Plotkin, 2016, 2017) (**Supplementary Figure 1**). Moreover, slower rates of environmental change are correlated with larger probabilities of fixation and larger probabilities of phenotypic memory that maximize the probability of fixation of the new allele (**Figure 4**). These results are intuitive and easy to interpret: it is beneficial for the *a* allele to have some phenotypic memory within each environment, as this helps the high-fitness realizations of the allele, while having little effect on its low-fitness realizations. However, too much phenotypic memory can be detrimental, because the *a* lineage will be “stuck” with a potentially deleterious phenotype for longer. The optimal amount of phenotypic memory is larger for slower-changing environments, as this allows the rates of phenotypic switching to be tuned to the environmental stochasticity and creates the optimal amount of phenotypic diversity for the given environmental rate of change (Carja et al., 2014b). For comparison, we include the probability of fixation for a neutral trait with no fitness advantage (horizontal lines, **Figure 4**). A phenotypically variable ?a? allele is advantageous over a wide range of phenotypic memory, and this advantage increases as the environmental rate of change increases.

**Figure 4:**
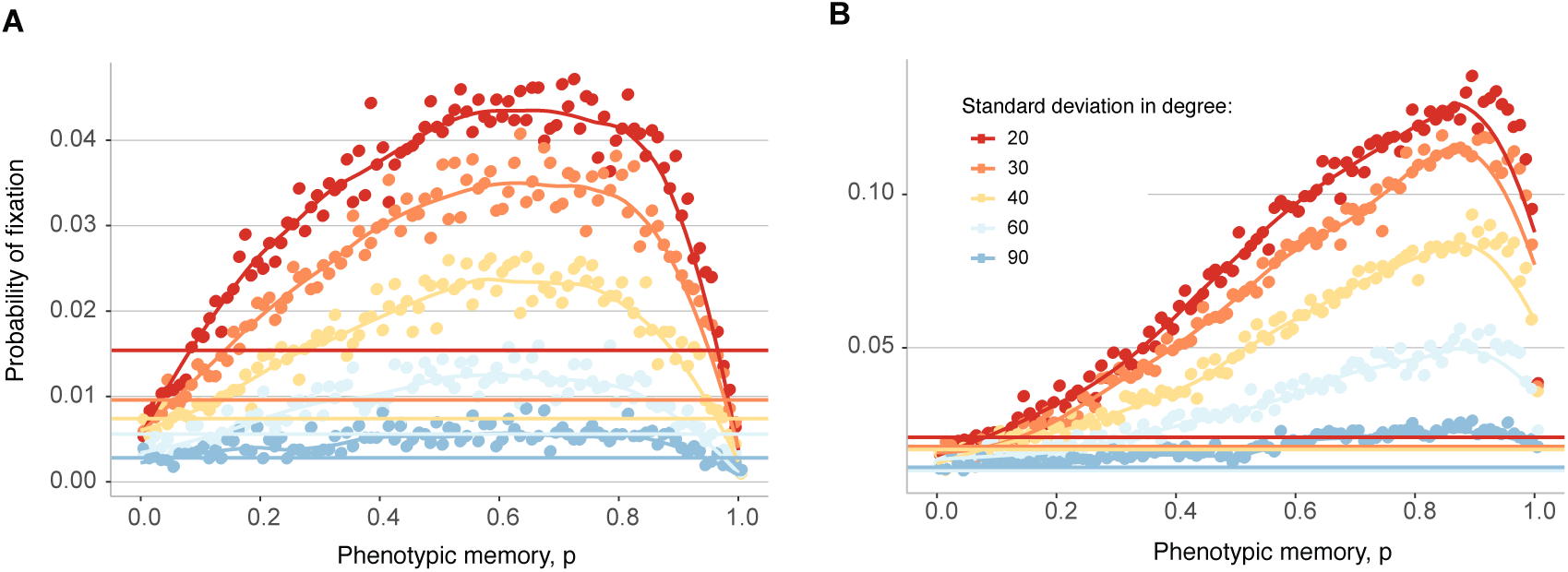
Probability of fixation for different values of the standard deviation in degree of the contact network. The lines are splines, while the dots represent the ensemble average across 5000 replicate Monte Carlo simulations. *N* = 500, Φ_*A*,*E*1_ = 0.8, Φ_*A*,*E*2_ = 0.6, Var(Φ_*a*_)=0.04, mean degree of the contact network is fixed at 40. **Panel A**: Environmental rate of change *n* =5. **Panel B**: Environmental rate of change *n* = 30. The horizontal lines represent the probability of fixation for a neutral allele entering the population, with colors as in the legend.

As a network grows and evolves, more connections are formed. In real-life networks, it has been observed that nodes that are already well connected can be more likely to acquire new connections, a concept known as ‘preferential attachment’ (Barabási and Albert, 1999; Dorogovtsev et al., 2000; Yook et al., 2002; Jeong et al., 2003). As the power of preferential attachment increases, new connections are more concentrated at well-connected nodes (the hubs of the network), and the standard deviation in degree of the network increases. We next focus on the differences in spatial population structure that result when networks have different levels of preferential attachment, and we assess the effect of these differences on the evolutionary advantage of a phenotypically plastic allele *a*. Specifically, we analyzed networks for which we tuned the power of preferential attachment while keeping the mean degree constant, which translated to changes in the standard deviation in degree.

We find that the fixation probability of a new phenotypically variable allele *a* depends highly on the heterogeneity of network structure, and that it can be markedly lower for networks with higher variance in degree, for a wide range of rates of environmental change (**Figure 4** and **Supplementary Figure 2**). This effect of the standard deviation in degree is shown to be much stronger than the effect of the mean degree of the contact network.

**Figure 5** shows that, for a given value of the probability of phenotypic memory *p*, (in this figure *p* = 0.5), the probability of fixation of the *a* allele is maximized for well-mixed populations and decreases as the standard deviation of degree is increased, across a wide range of rates of environmental stochasticity. This lower fixation probability for new mutations compared to well-mixed population points to slower adaptation as mutations become rare. Hence, population structure acts as a suppressor of selection, tilting the balance of forces towards drift and against selection and slowing down evolutionary dynamics.

**Figure 5:**
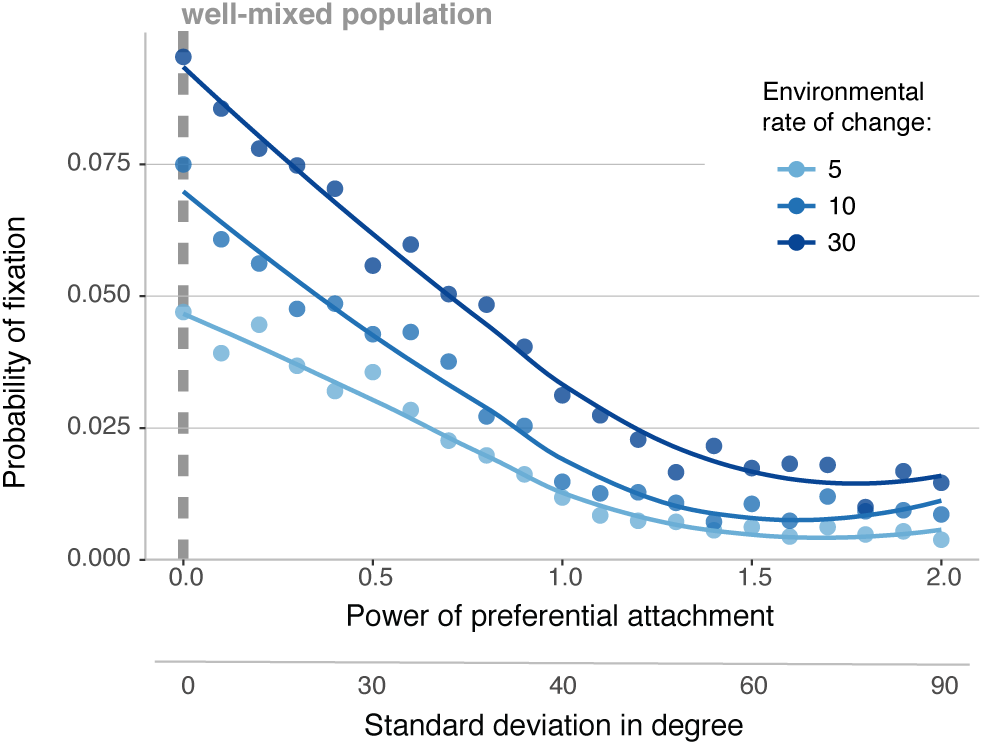
Probability of fixation of a new phenotypically plastic allele as a function of standard deviation of degree for the network for different rates of environmental change. The lines are splines, while the dots represent the ensemble average across 5000 replicate Monte Carlo simulations. *N* = 500, Φ_*A*,*E*1_ = 0.8, Φ_*A*,*E*2_ = 0.6, Var(Φ_*a*_)=0.04, mean degree of the contact network is fixed at 40. Here the phenotypic memory is fixed at *p* = 0.5. When the power of preferential attachment is 0, the results are representative of a well-mixed population. The *x*-axis shows the power of preferential attachment, and a secondary *x*-axis (below) shows the equivalent standard deviation in degree.

The intuition behind these results is simple. When a novel mutant arises in a random node of the network, it is much more likely to arise in a node of small degree for networks with high variance in degree (when the mean degree is kept constant). While the network does indeed contain bigger, higher-degree nodes, they are rare, and these hubs can only aid in the trait’s spread once they are reached. On average, however, when the power of preferential attachment is high the mutation is more likely to appear in nodes of small degree, and correlations in degree mean that these nodes of small degree are linked to other nodes of small degree. The architecture of this population structure constrains spread and increases the probability that the new mutation, even if beneficial, is lost from the population. The higher the variance, the stronger this effect, and the lower the average degree of most network nodes.

## Discussion

Human interactions are generally not random; they are often structured by geography, social networks, language, and other cultural factors. The aim of this study is to understand how network topology—in particular, heterogeneity in degree—shapes probabilities of fixation of phenotypically variable alleles, and thus to hint at how population structure shapes the rate of evolution in cultural systems. Studying the role of population structure is complicated by the fact that networks differ in many structural properties, and it is difficult to study one network feature independent of others. Degree distribution has been shown to be an essential characteristic of network structure (Maslov and Sneppen, 2002; Kossinets and Watts, 2006), and previous studies have identified network properties, such as individual variation in number of contacts, as an important determinant of disease spread (Newman, 2002; Bansal et al., 2007; Eames and Keeling, 2002; Shirley and Rushton, 2005; Salathé et al., 2010). In addition, heterogeneity in degree influenced the cascading spread of a neutral trait, such as a cultural fad, in a threshold model: increased heterogeneity in degree decreased the likelihood that such a fad would sweep through the population (Watts, 2002)

Throughout this paper, we have considered a model of cultural evolution taking into account two important aspects: the fact that cultural traits have partially heritable phenotypic variation and the fact that individuals’ interactions are not random in a population, but instead structured in local contact networks. We have modeled a single new mutation that can increase cultural phenotypic variability; we introduce this new plastic mutant at time *t* = 0 on a random node on the network and study its probability of fixation. This allows us to ask questions about the rate of cultural evolution in structured populations and interrogate whether spatial population structure in this case is an amplifier or a suppressor of selection. We have found that the probability that this new mutant fixes in the population is linked to the properties of the network that describes the populations’s interactions. When we introduce a new cultural bet-hedging mutant with multiple possible phenotypic states, the probability that this mutant spreads to fixation in the population increases when the individuals on the network have more connections (increased mean degree, **Figure 4**) but decreases when these connections are more unevenly distributed (increased standard deviation in degree, **Figure 5**). In other words, the network structure acts to inhibit the spread of bet-hedging mutations compared to well-mixed populations, slowing down the dynamics and tilting the evolutionary balance away from selection and towards drift. Further, the spread of this phenotypically plastic trait is particularly hindered by an uneven social network in which some individuals are well-connected hubs of information but many individuals have few connections.

The degree of cultural memory and the rate of environmental change also interact with network structure to produce novel cultural evolutionary patterns. Stronger cultural memory (*p*) increases the likelihood of fixation of the phenotypically plastic mutant allele, but only to a point. If the fidelity of cultural transmission from parent to offspring is too high, the high-fitness phenotype can spread, but after the environment changes the lower-fitness phenotype will then be overrepresented in the *a* population. Thus, there is an intermediate optimum level of cultural memory that depends on the environmental rate of change; slower rates of environmental change favor higher levels of cultural memory than faster rates do. These slower rates of environmental change are also correlated with larger probability of fixation of the mutant allele (**Figure 4**). Further, as the probability of fixation of the variable trait increases, it also spreads more quickly through the population (**Figure S3**), thus linking optimal levels of cultural memory to the rapid fixation of the phenotypically variable trait.

Such a model could be applied to many cultural systems; here we have given the example of hunting and gathering skills as cultural traits that can be fully or partially heritable and can be affected by the environment. In our example, individuals with the cultural allele *A* have a fixed food-source preference, such as gathering edible plants from the environment, and the culturally transmitted knowledge about which foods are edible. In contrast, individuals with the *a* allele can learn one of multiple hunting techniques that might involve specialized tools. Descendants of *a* individuals learn their parent’s hunting technique with probability *p*, which represents the degree of cultural memory. When the environment changes, food sources (such as specific plants or animals) that were previously abundant could become scarce, so preferences for those food sources could shift in their benefit to the individual.

Thus far, we have been considering environmental fluctuations as external changes that alter the fitnesses of the cultural traits in question. However, cultural traits can also alter their own environment, changing the selection pressures on both cultural and genetic traits, a process known as cultural niche construction (Laland et al., 2000; Odling-Smee et al., 2003; Creanza et al., 2012; Fogarty and Creanza, 2017). This is salient to our model: with the example of food-source choice as a cultural trait, we can envision that environmental factors such as rainfall or temperature can fluctuate over time, altering the fitness of the cultural practice of gathering or farming certain food sources. However, the cultural traits themselves can also act to facilitate changes to the environment. For example, gathering one type of food can lead to scarcity of that resource, reducing the fitness of utilizing that food source and increasing the fitness of shifting to another food source. Similarly, preferentially hunting large game might reduce their population sizes, possibly allowing the populations of smaller animals to increase and thus be a more abundant food source, increasing the fitness of other phenotypic variants of the hunting trait. From the perspective of food-producing societies, this process is also evident: specific crops use subsets of soil nutrients, altering the environment by depleting those resources and thus increasing the fitness of cultural traits favoring other crops. This underscores the importance of considering environmental fluctuations in models of cultural evolution, since culture itself can induce such fluctuations. Thus, for cultural traits, both external environmental changes and culture-induced niche construction prove to be crucial factors in evolutionary dynamics; in this model, we can represent both of these situations by implementing fluctuating environments.

We have based this model on a theoretical population-genetic framework, the death-birth Moran process, but when applied to cultural traits, it is not necessary to think of this process in literal terms of death and birth. In genetic models, an individual transmits its traits via reproduction; however, in the context of cultural evolution, this transmission can also occur between existing individuals. Thus, we can envision a Moran process for cultural traits such that, in every time step, an individual becomes culturally plastic and can learn from a neighbor. Thus, the death rate here can be conceptualized as a rate at which people become receptive to a new cultural model, at which point they could learn from a neighbor or sample from the distribution of phenotypes in the population. The process by which individuals reproduce in proportion to their fitness also has a cultural analogue: humans can show preferences for learning from individuals who have demonstrated success (Boyd and Richerson, 1988; Henrich and McElreath, 2003; McElreath et al., 2008; Mesoudi, 2011). Thus, in addition to representing vertical cultural transmission from parent to offspring, our model could also represent the spread of a behavior horizontally along a network of adults. For our hunting and gathering example, a randomly chosen individual in each time step could assess its neighbors to determine who has procured the most food recently and potentially copy the phenotype of that individual.

To understand how our results may generalize to other representations of the spread of cultural traits, we discuss some of implications of the model assumptions. First, by considering the evolution of one trait in a population, we must make the simplifying assumption that a cultural trait affects fitness equally across individuals. In reality, individuals have different cultural repertoires that would modulate the benefit of any given trait. We can easily see this in our hunting and gathering example: a simple additional trait—for example, the ability to fish, or the practice of sharing food among the group—might dampen an individual’s fitness benefit from specialized hunting skills. Further, a useful extension to the horizontal transmission framing of the model could be the assessment of one’s own fitness in comparison to surrounding individuals. Currently, when an individual is chosen to adopt a new trait, it will do so even when all of the surrounding individuals have a lower-fitness variant. This has been observed in cultural systems: conformist-biased cultural transmission can lead to the propagation of common cultural traits even when they are maladaptive or costly (Henrich, 2000). However, humans are also adept at assessing the prestige or success of an individual before deciding to learn from that individual, cultural phenomena termed prestige bias and success bias (Barnett and Pontikes, 2008; Reyes-Garcia et al., 2008); this could be modeled in our framework if individuals assessed the fitness of their potential tutors before deciding whether to alter their cultural trait. Success bias in horizontal transmission could thus potentially aid the spread of the phenotypically plastic allele, since the high-fitness forms of the trait would preferentially spread in a given environment.

In addition, we have modeled population structure as a static, fixed network; interactions between individuals in many real-world situations, however, are dynamic. For example, humans are mobile and their likelihood of interactions with other individuals can depend on their current geographic location. However, connections between certain individuals, such as genetic relatives, might be more stable over time. This modeling framework could usefully be applied both to real-world networks constructed from human interactions and to dynamic networks in which connections can form and break over time. Such rewiring of the network may dampen the effect of population structure if it alters the instantaneous degree distribution or maintains the degree distribution but changes which nodes are well connected. Finally, as with previous work in (epi)genetic systems, these models focus on a non-random, periodic environmental fluctuation (but see (Carja et al., 2014b)). There is considerable scope for extensions of this model to account for random fluctuations in environment as well as fluctuations with more than two possible environmental states.

In conclusion, we show that network structure, cultural plasticity, and environmental fluctuations interact with one another to produce complex cultural evolutionary dynamics. In particular, heterogeneity in contact structure suppresses cultural evolution of a phenotypically varying trait by lowering the fixation probability a newly arising plastic mutation. This finding is surprising because social network research in other fields, such as marketing, indicates that hubs in a network could have an outsized impact on facilitating the spread of information (Goldenberg et al., 2009). These contrasting results might suggest a spectrum of influence for network hubs depending on the type of cultural transmission in a system, warranting further investigation. For example, certain cultural traits might be easily observed and adopted without close interpersonal interactions, facilitating one-to-many transmission and augmenting the importance of network hubs. In contrast, if other cultural traits are primarily passed through parent-to-offspring transmission or through extended teaching, then network hubs might be far less influential in, or even detrimental to, the spread of information.

## Acknowledgments

The authors would like to thank Kate Snyder, Abigail Searfoss, Cristina Robinson, Parker Rundstrom and members of the Creanza Laboratory for useful feedback and suggestions. This research was done using resources provided by the Open Science Grid, which is supported by the National Science Foundation award 1148698, and the U.S. Department of Energy’s Office of Science.

## List of Figures

**Figure S1:**
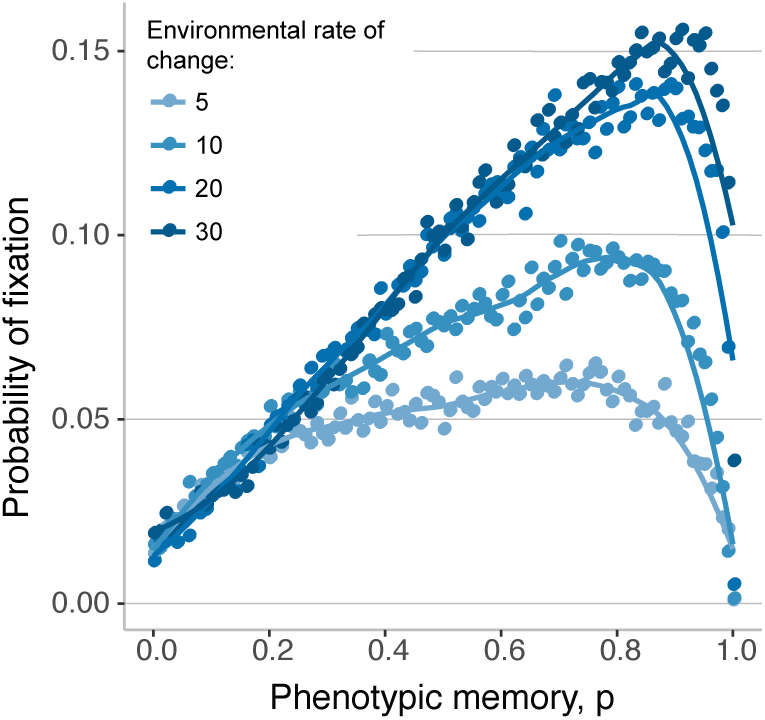
Probability of fixation of the *a* allele for well-mixed populations. The lines are splines, while the dots represent the ensemble average across 5000 replicate Monte Carlo simulations. Here *N* = 500, Φ,_*A*,*E*1_ = 0.8, Φ_*A*,*E*2_ = 0.6, Var(Φ_*a*_)=0.04 and the rate of environmental changes as presented in the legend.

**Figure S2:**
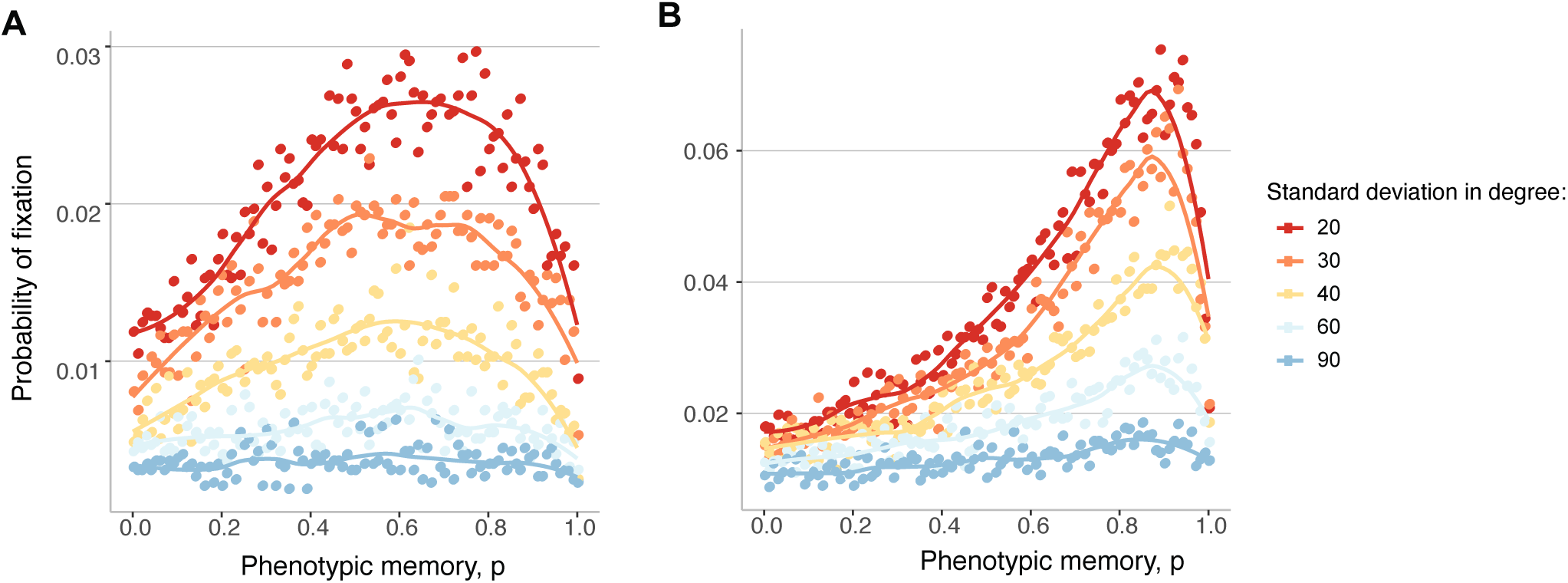
Probability of fixation for different values of the standard deviation in degree of the contact network. The lines are splines, while the dots represent the ensemble average across 5000 replicate Monte Carlo simulations. Here *N* = 500, Φ_*AE*1_ = 0.8, Φ_*A*,*E*2_, = 0.6, Var(Φ_*a*_)=0.01, mean degree of the contact network is fixed at 40. **Panel A**: Environmental rate of change *n* = 5. **Panel B**: Environmental rate of change *n* = 30.

**Figure S3:**
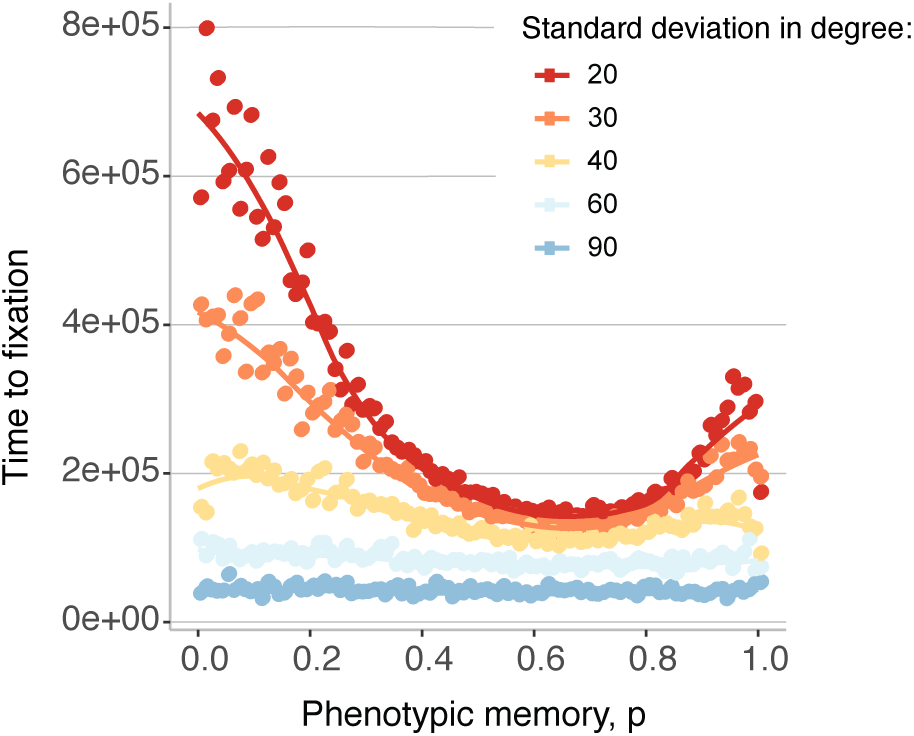
Time to fixation for different values of the standard deviation in degree of the contact network. The lines are splines, while the dots represent the ensemble average across 5000 replicate Monte Carlo simulations. Here *N* = 500, Φ_*A*,*E*1_ = 0.8, Φ_*A*,*E*2_ = 0.6, Var(Φ_*a*_)=0.04, mean degree of the contact network is fixed at 40. Environmental rate of change *n* = 5.

**Table 1:** Other network features for the degree distributions used in the manuscript.

| Mean degree | Standard deviation in degree | Average path length | Clustering Coefficient |
| --- | --- | --- | --- |
| 40 | 20 | 2.2 | 0.15 |
| 40 | 30 | 2.2 | 0.17 |
| 40 | 40 | 2.2 | 0.18 |
| 40 | 60 | 1.8 | 0.16 |
| 40 | 90 | 1 | 0.1 |

